# CAVE: Connectome Annotation Versioning Engine

**DOI:** 10.1101/2023.07.26.550598

**Authors:** Sven Dorkenwald, Casey M. Schneider-Mizell, Derrick Brittain, Akhilesh Halageri, Chris Jordan, Nico Kemnitz, Manual A. Castro, William Silversmith, Jeremy Maitin-Shephard, Jakob Troidl, Hanspeter Pfister, Valentin Gillet, Daniel Xenes, J. Alexander Bae, Agnes L. Bodor, JoAnn Buchanan, Daniel J. Bumbarger, Leila Elabbady, Zhen Jia, Daniel Kapner, Sam Kinn, Kisuk Lee, Kai Li, Ran Lu, Thomas Macrina, Gayathri Mahalingam, Eric Mitchell, Shanka Subhra Mondal, Shang Mu, Barak Nehoran, Sergiy Popovych, Marc Takeno, Russel Torres, Nicholas L. Turner, William Wong, Jingpeng Wu, Wenjing Yin, Szi-chieh Yu, R. Clay Reid, Nuno Maçarico da Costa, H. Sebastian Seung, Forrest Collman

## Abstract

Advances in Electron Microscopy, image segmentation and computational infrastructure have given rise to large-scale and richly annotated connectomic datasets which are increasingly shared across communities. To enable collaboration, users need to be able to concurrently create new annotations and correct errors in the automated segmentation by proofreading. In large datasets, every proofreading edit relabels cell identities of millions of voxels and thousands of annotations like synapses. For analysis, users require immediate and reproducible access to this constantly changing and expanding data landscape. Here, we present the Connectome Annotation Versioning Engine (CAVE), a computational infrastructure for immediate and reproducible connectome analysis in up-to petascale datasets (∼1mm^3^) while proofreading and annotating is ongoing. For segmentation, CAVE provides a distributed proofreading infrastructure for continuous versioning of large reconstructions. Annotations in CAVE are defined by locations such that they can be quickly assigned to the underlying segment which enables fast analysis queries of CAVE’s data for arbitrary time points. CAVE supports schematized, extensible annotations, so that researchers can readily design novel annotation types. CAVE is already used for many connectomics datasets, including the largest datasets available to date.

## Introduction

Volume Electron Microscopy (EM)^1, 2^ provides an exquisite view into the structure of neural circuitry and is currently the only technique capable of reconstructing all synaptic connections in a block of brain tissue. EM imagery not only facilitates the reconstruction of neuronal circuits but also enables scientists to combine^3–8^ them with rich ultrastructure visible in these images (Fig. 1a). An increasing set of ultrastructural features, such as synapses^3, 4, 9, 10^, their neurotransmitter identity^4, 11^, and mitochondria^3–7^, can be extracted automatically through machine learning methods. In addition, human experts have long used electron microscopy to make a rich set of observations about cellular and sub-cellular processes, including the localization of a wide range of organelles and cell-to-cell interactions^12–15^. When combined with neuronal reconstructions, these datasets enable new analyses of richly annotated connectomes^16–22^. This trend is mirrored in other data-intensive fields such as genome sequencing^23^ and large scale astronomy surveys^24^, where raw data is iteratively enriched as increasingly accurate and diverse sets of annotations are added.

**Figure 1.**
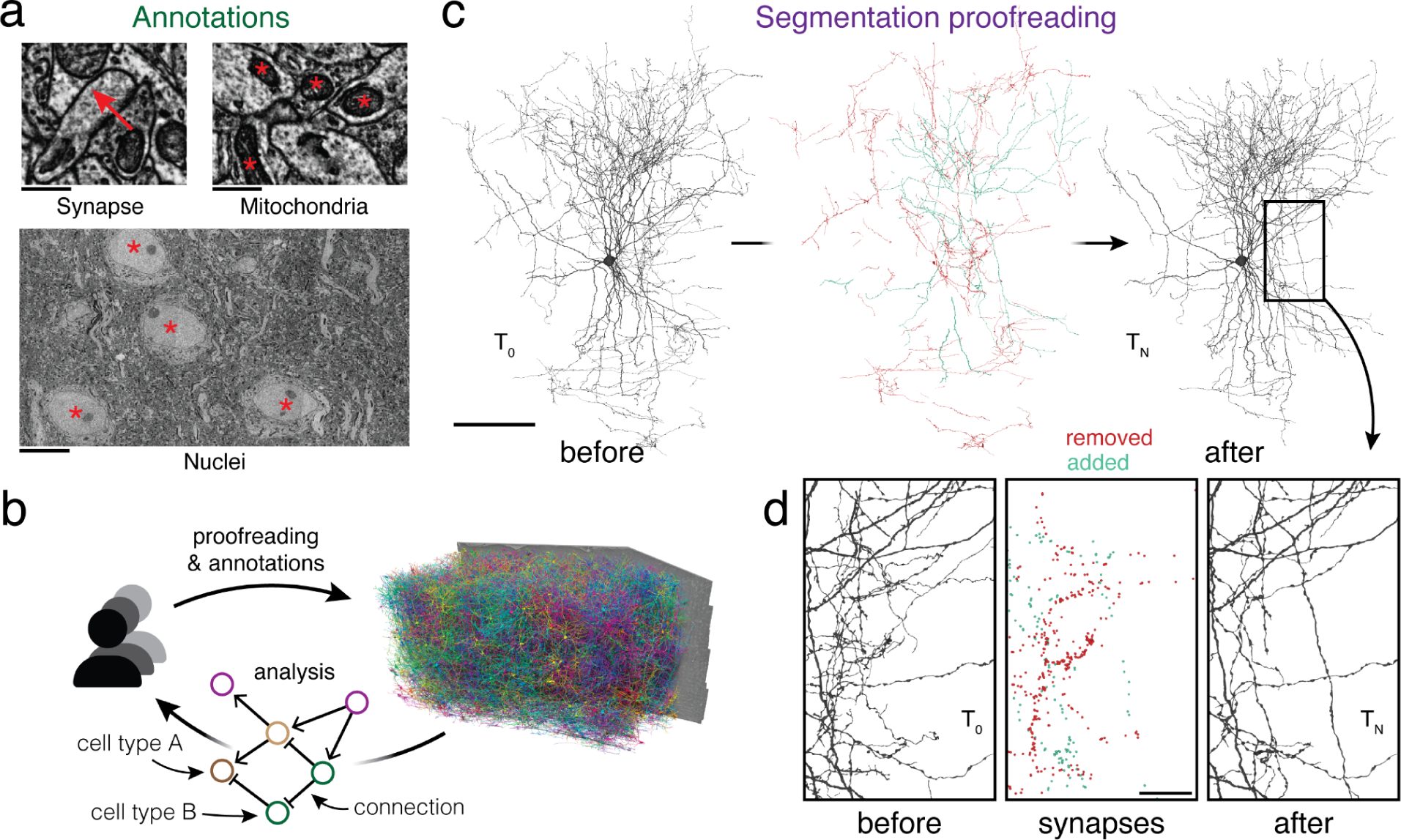
Proofreading and analysis of connectomics datasets. (a) A rich set of ultrastructural features can be extracted from EM images and used for analysis. The corresponding ultrastructural features are annotated with a red *. The synapse is annotated with a red arrow pointing from the pre- to the postsynaptic site. (b) Large connectomics datasets are proofread, annotated, and analyzed by a distributed pool of users in parallel. (c) Proofreading adds and removes fragments from cell fragments (left: before proofreading, center: removed and added fragments, right: after proofreading). (d) Synapse assignments have to be updated with proofreading. All synapses (within the cutout) that were added and removed though the proofreading process of the cell in (c) are shown. Scale bars: 100 µm (c), 1 µm (a: synapse, mitochondria), 10 µm (a: nuclei), 20 µm (d)

Today, neurons in EM datasets are extracted through automated approaches^25–29^ in order to scale analyses to increasingly larger volumes^16, 18–20^. However, manual proofreading of automated segments is still necessary to achieve reconstructions suited for analysis^30^.

Proofreading of large datasets takes years, but even partial connectomic reconstructions produced along the way are useful for analysis. This raises the need for software infrastructure that facilitate concurrent proofreading within large collaborations or entire communities of scientists and proofreaders each working on their individual analyses (Fig. 1b). However, existing tools and workflows only support static exports after proofreading was completed^31, 32^.

To enable this shift to collaborative proofreading and analysis of connectomics datasets, we created the Connectome Annotation Versioning Engine (CAVE). CAVE introduces a set of new methods for connectome analysis and combines them to a coherent system. For proofreading, CAVE builds on the ChunkedGraph^33, 34^. Like previous systems^31, 35–39^, the ChunkedGraph represents cells as connected components in a supervoxel (groups of voxels) graph. It is currently the only system for neuron-based proofreading by a distributed community but was too costly to be used on petascale datasets (∼1mm^3^ of brain tissue). Here, as one part of CAVE, we introduce the next generation of this system, the ChunkedGraph v2, which scales proofreading to petascale dataset through a more cost-efficient storage implementation.

Ongoing proofreading presents a challenge for analysis: edits not only add and remove fragments from cell segments (Fig. 1c), but they also change the assignment of cell labels and ultrastructural features such as synapses (Fig. 1d), and require recalculations of morphological neuron features (e.g. volume and area) and neuronal representations (e.g., skeletons). Previous systems that combined reconstruction, annotation and analysis^40–43^ only supported manual cell reconstructions and manual annotations.

As a second part of CAVE, we addressed the challenge of supporting analysis of proofreadable cell segmentations in conjunction with annotations produced by automated methods and individual users. CAVE enables fast computation of morphological neuron features and representations at any time, including immediately after an edit, through an extension to the ChunkedGraph. We introduce a new scheme for storing annotations which binds annotations to segment IDs at specific points in time in a process we call “materialization.” We show that CAVE’s annotation and proofreading systems support fast queries of the data for any point in time by combining traditional database queries with ChunkedGraph-based tracking of neuron edit histories. This enables CAVE to answer analysis queries with no delays after an edit, as well as queries of the state of data at arbitrary timepoints.

Taken together, CAVE manages concurrent proofreading, annotation, and annotation assignment while offering queries to the data for any point in time to support flexible and reproducible analysis by a distributed group of users. CAVE is already used to host 5 published datasets where it tracks almost 2 billion annotations and has recorded over 4 million edits by over 500 unique users from across the globe. CAVE facilitated the reconstruction of the first whole-brain adult connectome with FlyWire^20, 44^, supports the FANC community reconstructing the Drosophila VNC^45^, and hosts the cubic millimeter-scale MICrONS volumes^18^. Currently, CAVE is the only system to support concurrent analysis, annotation and proofreading by a distributed community, and in particular proofreading at the cubic millimeter-scale.

## Results

### Real-time collaborative proofreading of petascale reconstructions

In a perfect segmentation, all voxels (3D pixels) within the same cell are labeled with the same ID (Fig. 2a). Here, we refer to a group of voxels with the same ID as a “segment”. In an automated segmentation, most segments require subsequent proofreading to create accurate neuron reconstructions. Here, we use the term “proofreading” exclusively for edits to the segmentation but CAVE supports editing of annotations as well. Proofreading an automated segmentation is 10-100x faster than purely manual reconstruction^20, 30, 39, 46, 47^, but proofreading of a large dataset may still go on for years. An ideal system should allow real-time collaboration by many proofreaders, including both humans and machines, and make the results available for concurrent analysis and discovery efforts.

**Figure 2.**
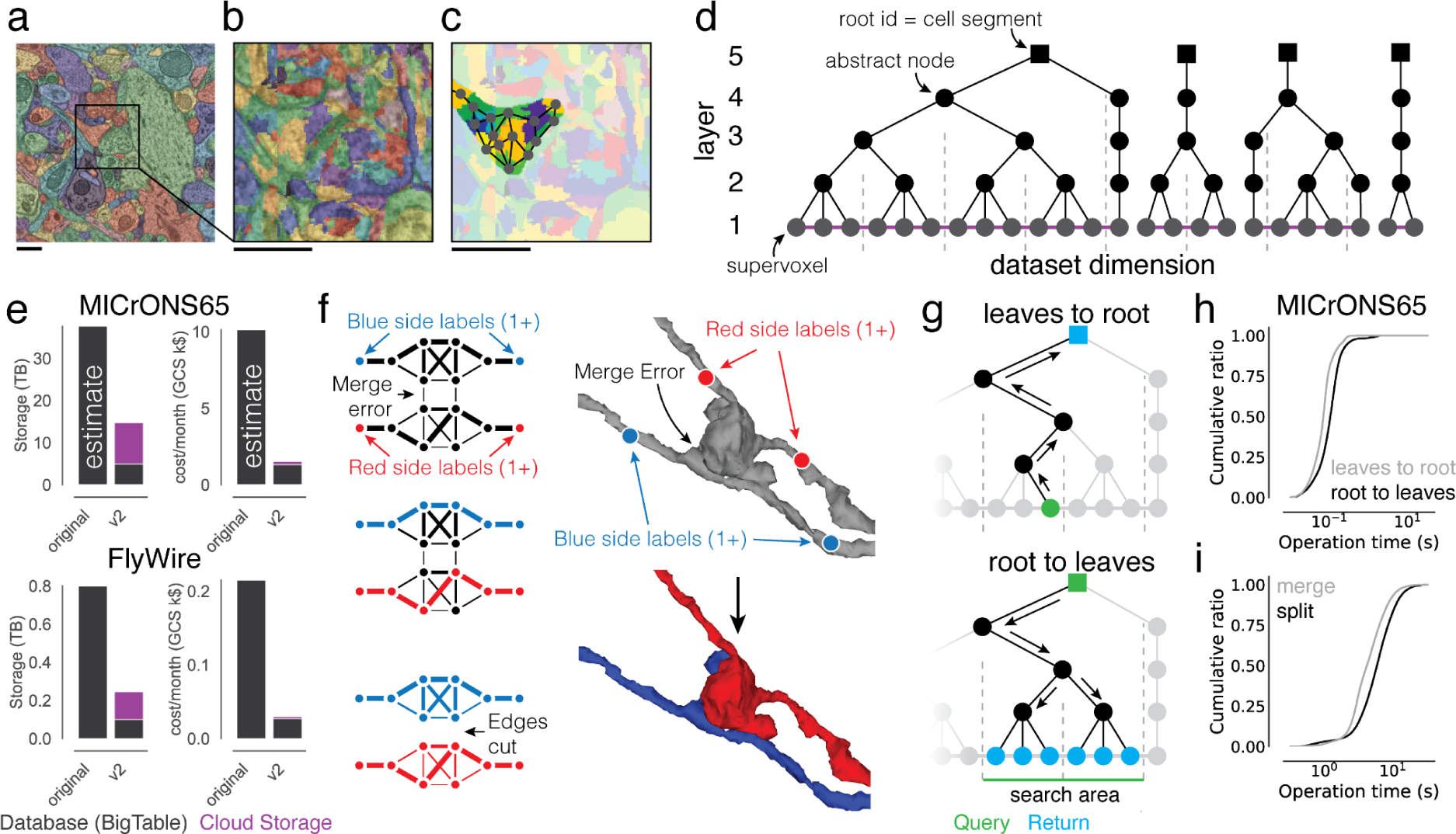
Scaling the ChunkedGraph to petascale datasets. (a) Automated segmentation overlaid on EM data. Each color represents an individual putative cell. (b) Different colors represent supervoxels that make up putative cells. (c) Supervoxels belonging to a particular neuron, with an overlaid cartoon of its supervoxel graph. This panel corresponds to the framed square in (a) and the full panel in (b). (d) One-dimensional representation of the supervoxel graph. The ChunkedGraph data structure adds an octree structure to the graph to store the connected component information. Each abstract node (black nodes in levels >1) represents the connected component in the spatially underlying graph. (e) Storage and costs for the supervoxel graph storage under the original and the improved implementation (v2). (f) To submit a split operation users place labels for each side of the split (right top). The backend system first connects each set of labels on each side by identifying supervoxels between them in the graph (left). The extended sets are used to identify the edges needed to be cut with a max-flow min-cut algorithm. (g) Examples of graph traversals for looking up the root id for a supervoxel id (top) and supervoxel ids for a root id within a spatially defined search area (bottom). Note that only part of the graph needs to be traversed. (h) Performance measurement from real-world user interactions measured on the ChunkedGraph server for different types of reads and (i) edits. Scale bar: 500 nm

**Extended Data Figure 2-1.**
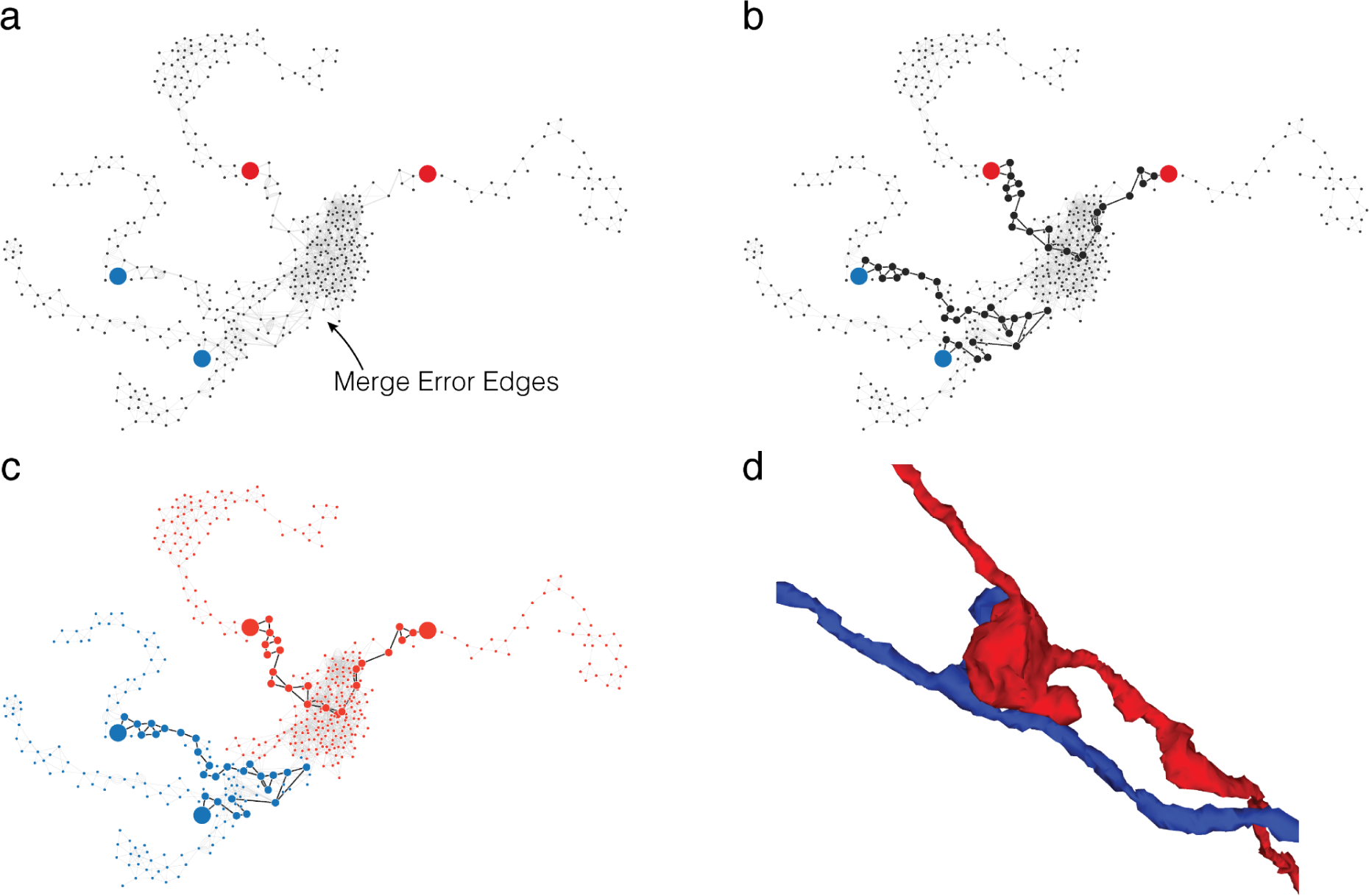
Translating user inputs to graph splits. (a) Bipartite split labels are applied to locations in space. (b) The closest supervoxels to label points are identified (red/blue dots). The supervoxel graph in the neighborhood of the labeled points is computed (graph), weighted by affinity between supervoxels. (c) Vertices along the shortest paths between each pair of red/blue labels are found (black dots and edges). Backup methods prevent overlap between paths. (d) Affinity between vertices along shortest paths is set to infinity and min cut is computed on the path-augmented supervoxel graph.

**Extended Data Figure 2-2.**
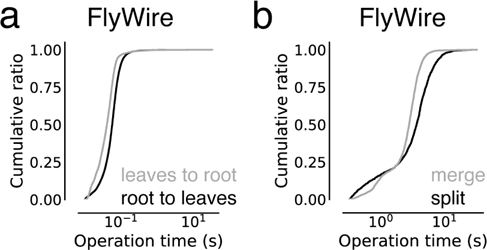
ChunkedGraph performance measurements on FlyWire. These measurements are from the improved ChunkedGraph implementation using the same FlyWire supervoxel graph that was used for the original implementation^34^. (a) Performance measurement from real-world user interactions measured on the ChunkedGraph server for reads, specifically leaves to root (median=41ms, N=13,410) and root leaves (median=55ms, N=50,001) operations, and (i) edits, specifically merge (median=2,734ms, N=4,189) and split (median=3,486ms, N=2,875) operations.

These requirements are met by the ChunkedGraph proofreading system, whose design was described previously ^33, 34^. Like previous proofreading systems^31, 35, 36^, the ChunkedGraph stores the segmentation as a graph of atomic segments, called supervoxels (Fig. 2a,b). Connected components in this graph represent neurons (Fig. 2c). The ChunkedGraph introduced a new representation of the segmentation as a spatially chunked hierarchical graph of supervoxels (Fig. 2d) where root nodes are individual cell segments, and leaf nodes are supervoxels. To achieve high performance, the ChunkedGraph requires a database featuring low-latency random row reads such as BigTable^48^ which can add significant cost to its deployment. CAVE uses the ChunkedGraph as proofreading backend and hosts it as a cloud service for world-wide access. Here, we describe two significant advancements to the ChunkedGraph to make it viable for petascale datasets.

First, we reimplemented the ChunkedGraph creating the ”ChunkedGraph v2”. The initial ChunkedGraph version targeted proofreading of the FlyWire^34^ and the MICrONS phase 1 dataset (https://www.microns-explorer.org/phase1)^17,33,49^ datasets but turned out to be prohibitively costly for 50-100x larger petascale datasets like MICrONS65 (Fig. 2e). We redesigned the storage to a hybrid scheme in which supervoxel edges (purple lines in Fig. 2d), which are only needed for edits, are compressed and stored on conventional storage while the octree hierarchy remains stored in BigTable (Fig. 2e, number of supervoxels in MICrONS65: 112 billion). The reimplemented ChunkedGraph v2 reduced cost by >6.5x, low enough to support proofreading at the petascale (Fig. 2e).

Second, we improved how user edits are processed to speed up proofreading. To make proofreading accessible, proofreaders should not need to be aware of the underlying data structures. Instead, users perform edits by placing line connectors for merges and points for splits (Fig. 2f). The ChunkedGraph implements splits with a max-flow min-cut operation where user-selected supervoxels are labeled as sources and sinks to find the edges in the graph that should be removed. To aid split operations, we implemented an algorithm that uses a small number of locations coarsely surrounding the merge error, making the resulting topology of the split robust to the precise location or number of labels (Fig. 2f, Ext. Data Fig. 2-1). To speed up merging of many fragments, we added a multi-merge operation to neuroglancer allowing users to execute merge operations in parallel.

We measured view and edit performances during real-world proofreading of the ChunkedGraph v2 on the MICrONS65 dataset, the largest currently available, and the FlyWire dataset which we could use for comparison with the original implementation. For viewing the segmentation, the user selects a supervoxel by clicking a location in space, and the system retrieves all supervoxels that belong to the same segment within the field of view (Fig. 2f) in two steps. First, the ChunkedGraph is traversed from the selected supervoxel to the root node (Fig. 2f top). For MICrONS65, the ChunkedGraph v2 responded with a median time of 70.3 ms and 95th percentile of 247 ms (server-side performance, N=94,052) (Fig. 2g). Second, the search proceeds down the hierarchy to retrieve all supervoxels within a bounding box around the user’s field of view (Fig. 2f bottom). Here, the ChunkedGraph leverages the octree structure to avoid the retrieval of supervoxels out of the user’s view. We measured median response times of 104 ms and a 95th percentile of 291 ms (batched requests for multiple segments, N=182,411). Next, we tested edit operations. The server completed merge operations with a median time of 4,114 ms (N=25,839) (Fig. 2h) and splits with a median edit completion time of 5,810 ms (server-side, including the logic to identify sources and sinks; N=21,889) (Fig. 2h). Repeating this analysis with the FlyWire dataset show that viewing operations performed equally for the ChunkedGraph v2 and the original version while edit operations showed a modest slow down of ∼1.5s (Ext. Data Fig. 2-2). Notably, the performance for edits was only ∼1.6x times slower on the MICrONS65 dataset, even though it is 67x larger than FlyWire, illustrating the scalability of the ChunkedGraph system.

### Morphological analysis of proofread neurons

Proofreading is often driven by specific analysis goals. Being able to analyze cells as they are being corrected is important for analysis and to guide further proofreading. For instance, morphological information about a cell, e.g. volume and area, as well as morphological representations such as skeletons and meshes are used in many analyses. Skeletons are sparse representations of neurons that have proven useful for analysis and matching of neurons between datasets, including datasets of different modalities^50–52^. Computing these measurements and representations usually requires loading the entire segmentation of a cell which can span a significant part of a dataset. Recomputing these features from scratch after every edit is prohibitively time consuming and costly.

We leveraged the ChunkedGraph tree structure to cache and reuse meshes and morphological features for spatial chunks, and only recompute features in the regions of a cell that changed due to an edit. This L2-Cache (Fig. 3a), named after its use of level 2 of the ChunkedGraph hierarchy, is populated automatically through a queuing system (here: Pub/Sub) system after every edit. Every edit produces a list of new level 2 nodes and associated level 2 IDs (L2-IDs), for which a scalable microservice computes new meshes and a set of features, e.g. volume, area, representative coordinate, PCA components (see Methods).

**Figure 3.**
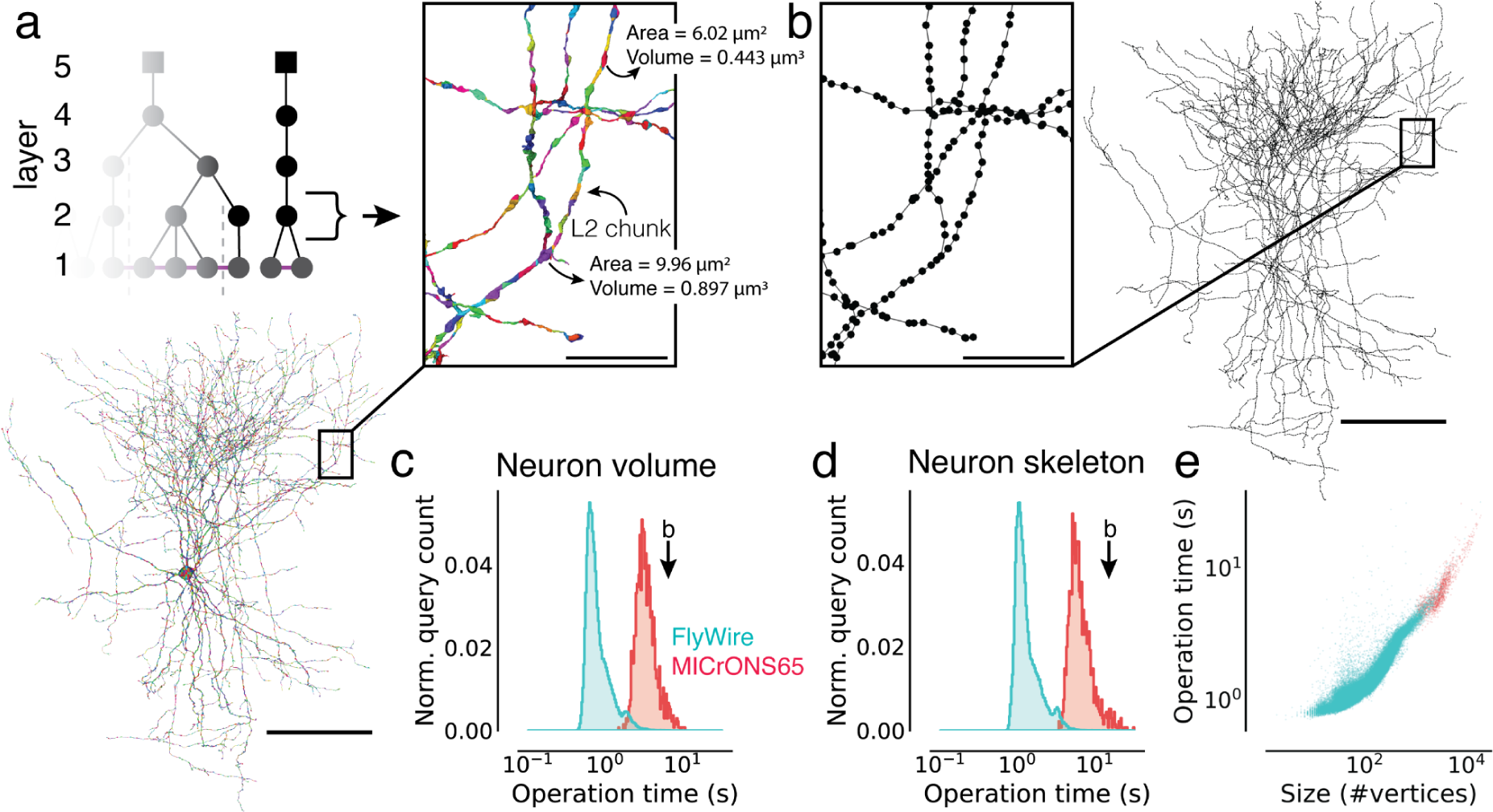
Fast calculation of morphological features and skeletons. (a) The basket cell from Fig. 1c broken into L2 chunks where each chunk is colored differently. For each chunk, the L2-Cache stores a number of features such as area, volume, and representative coordinate. (b) A skeleton derived from the ChunkedGraph and L2-Cache without consulting the segmentation data. (c) Client-side timings for calculating neuron volumes using the ChunkedGraph and the L2-Cache for neurons in FlyWire and MICrONS65. The timing for the neuron in (b) is highlighted. (d) Client-side timings for creating skeletons from the ChunkedGraph and the L2-Cache. (e) Client-side timings for creating skeletons plotted against the size of the skeletons. Each dot is a query for a single neuron. Scale bars: 100 µm, insets: 5 µm Combined with a fast retrieval of all L2-IDs belonging to a neuron (Ext. Data Fig. 3-1a), morphological features can be computed quickly. For instance, volume information can be computed within a median client-side time of 710 ms for FlyWire neurons and 3,176 ms for neurons in MICrONS65. The longer times for MICrONS65 can be explained by the larger size of the neurons (Ext. Data Fig. 3-1a-c).

To produce skeletons, the ChunkedGraph exposes a graph between L2-IDs, the L2-graph, which, when combined with locally computed representative coordinates from the L2-Cache, allow for rapid production of topologically correct skeletons. We implemented a graph based generalization of the TEASAR skeletonization algorithm ^53^ on the L2-graph to remove short and artificial branches introduced by the L2 chunk boundaries. Skeleton calculations of neurons in MICrONS65 and FlyWire took a median of 5,996 ms and 1,171 ms respectively with differences again being explained by the difference in size (Ext. Data Fig. 3-1d).

**Extended Data Figure 3-1.**
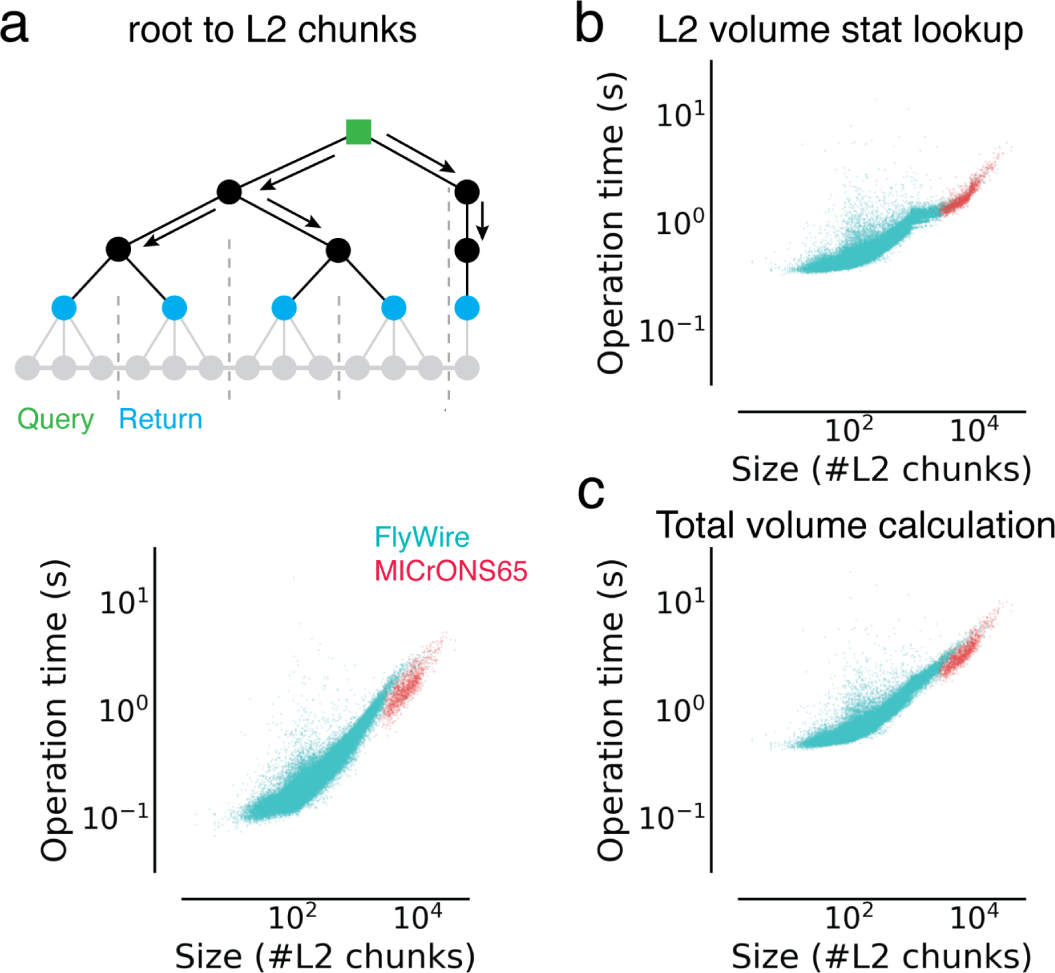
Analysis of timings to calculate morphological features. Each dot is a query for a single neuron. (a) Times to retrieve a list of L2 chunks for a neuron (root id). (b) Time to look up volume measurements for all L2 chunks belonging to a given neuron. (c) Total time to calculate volumes for neurons.

### Annotation schemes for rapid analysis queries

CAVE supports a diverse set of annotations from manual and automated sources by using flexible annotation schemas with a generic workflow engine. Every annotation is based on points in space that serve as spatial anchors and are accompanied by a set of metadata entries (Fig. 4c). Users can create new schemas that fit specific needs with arbitrary numbers of points (≥1) and metadata. To associate annotations with segments, the spatial points are bound to the underlying supervoxels (Fig. 4d) which can then be mapped to their associated root segment for any point in time using the ChunkedGraph. For instance, schemas describing synapses between two neurons contain two “bound spatial points’’ which are associated with the pre- and postsynaptic segments but vary in their additional parameters (e.g. size, neurotransmitter). In addition, reference schemas can be defined. Reference annotations are associated with annotations in another table via foreign key constraints (SQL) and can be used to track additional metadata without the need to create a copy of the annotation table itself.

**Figure 4.**
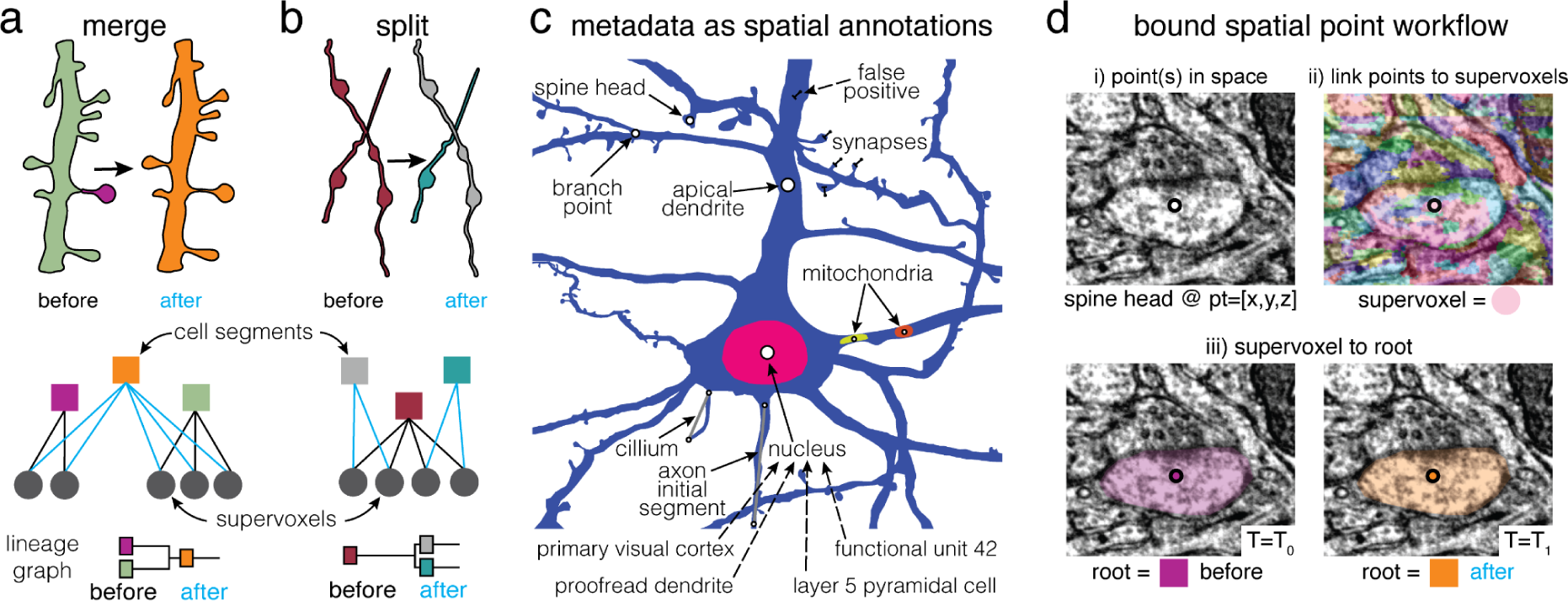
Annotations for proofreadable datasets. Basic operations of proofreading: (a) merging two objects and (b) splitting two objects. Each edit creates one or more new root objects (cell segments) that represent connected components of the supervoxel graph (octree levels not shown). The changes are tracked in a lineage graph of the altered roots. (c) Spatial points can be used to capture a huge diversity of biological metadata generated by either human annotators or machine algorithms. In CAVE annotations can be created as reference annotations which add additional metadata to existing locations (illustrated as dashed lines). (d) The annotation services handle all annotations through a generic workflow that depends only on expressing all annotations as collections of spatial points and associated metadata. (i) Spatial annotations mark the location of a feature such as a spine head. (ii) The materialization service retrieves the supervoxel id underlying all spatial points. (iii) This enables the materialization service to lookup the root id underneath that points at any given moment in time using the ChunkedGraph.

In comparison to other tools ^32^, CAVE is designed to be flexible about how users define their annotations so long as they include a bound spatial point. This allows the system to capture an expanding set of rich observations about the dataset, from small ultrastructural details, to observations about cell types and their anatomical locations. In fact, across MICrONS65^18^, MICrONS phase 1^17, 33, 49^, FlyWire^34^, and FANC^45, 54^, users have created over 120 annotation tables (including 29 reference tables), using 21 distinct schemas, capturing over 1.8 billion annotations. This includes tables marking synapse detections, reference annotations on those detections, nucleus locations, 62 distinct cell type tables, proofreading statuses, mitochondrial locations and functional co-registration points. Some tables are associated with individual studies making it easy to share data and reproduce analyses. We expect the diversity of observations to grow more rich over time and as further secondary analyses are performed.

CAVE maintains a “live” SQL database of all annotations. Users create annotation tables with any schema to which they add, remove and update individual annotations. Every time an annotation is added or updated, supervoxels underlying any bound spatial points are automatically retrieved. A service then frequently (e.g. 1/hour) updates the associated root segments of all annotations using the ChunkedGraph, which provides a list of updated root segments since the last database update (Fig. 4a,b). Infrequent copies of this database are produced and serve as materialized analysis snapshots. Due to the updates the live database is not suited for analysis queries as we cannot guarantee consistent versioning for multiple queries.

### Queries for arbitrary timepoints

Data analyses often require multiple queries to different annotation tables and use filters to reduce the data in the database to a manageable subset. A common example is the combination of a cell type query with a synapse query for a subset of neurons. This is usually achieved by filtering annotations with a set of root segments of interest. Because proofreading constantly changes the assignments of annotations to segments (Fig. 5a), all queries for one analysis need to be performed with the same version of the data, i.e. the same timestamp, to guarantee consistency and reproducibility. One way to ensure consistent queries to all tables is to query the materialized analysis snapshots (Fig. 5b). However, this limits a user’s ability to query the data immediately after fixing a segmentation error, a common scenario when doing exploratory analysis and proofreading in the dataset. This makes the snapshot system unsuitable for tracking proofreading, and requires a large number of snapshots to be kept to support continued analysis of past time points.

**Figure 5.**
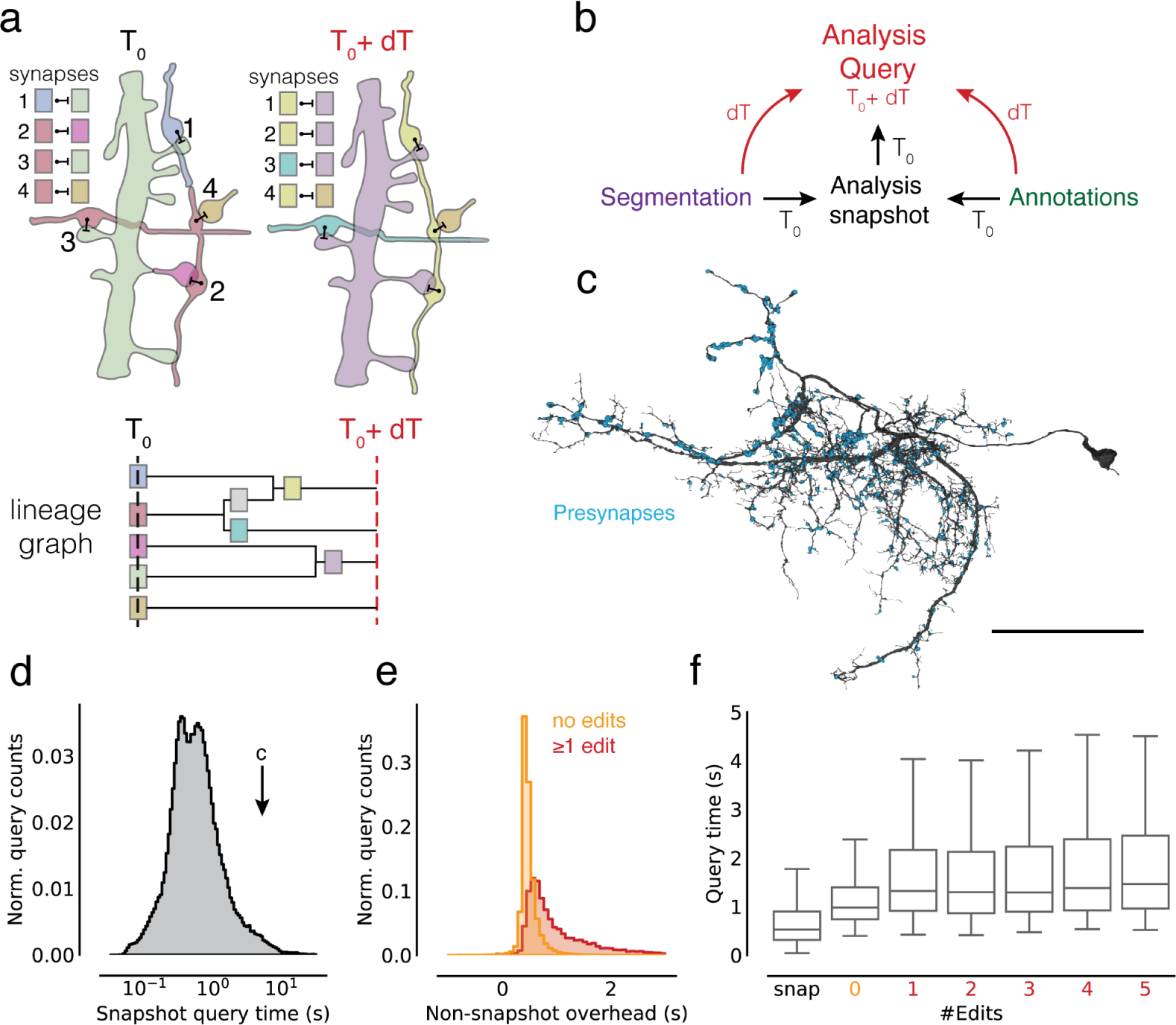
Querying the dataset for any time point. (a) Edits change the assignment of synapses to segment IDs. The lineage graph shows the valid IDs (colors) for each point in time. (b) Analysis queries are not necessarily aligned to exported snapshots. Queries for other time points are supported by on-the-fly delta updates from both the annotations and segmentation through the use of the lineage graph. (c) A neuron in FlyWire with all its automatically detected presynapses. (d) Time measurements for snapshot aligned queries of presynapses for one proofread neuron in FlyWire. (e) The difference between the snapshot and non-snapshot aligned presynapse queries. The two distributions differentiate cases without any edits to the queried neurons and cases with at least one edit to the queried neuron. (f) Presynapse query times for snapshot and non-snapshot aligned queries including cases where neurons were proofread with multiple edits. Scale bar: 50 µm

**Extended Data Figure 5-1.**
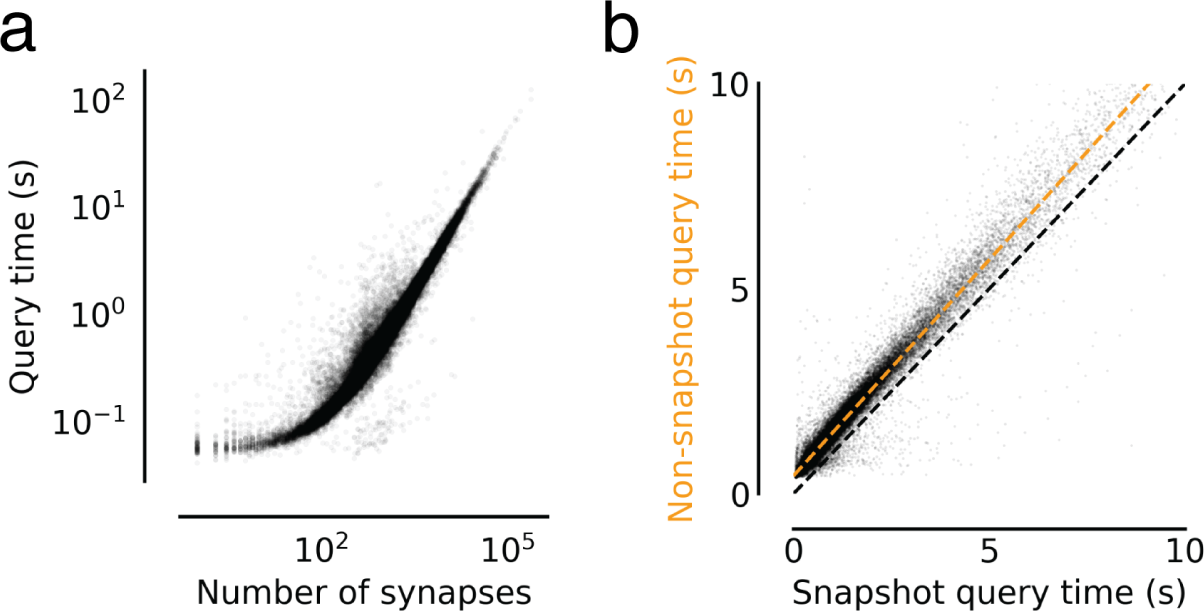
Annotation query timing analysis. (a) Query times from Fig. 5d versus the size of the query in number of presynapses. (b) Comparing snapshot and non-snapshot aligned presynapse queries for cases where the neuron was not edited between the snapshot and the query time. The difference is the overhead of the mapping logic. The green dashed line is a linear fit with intercept 0.44s and a slope of 1.05.

CAVE combines materialized snapshots with ChunkedGraph-based tracking of neuron edit histories to facilitate analysis queries for arbitrary time points (Fig. 5b). The ChunkedGraph tracks the lineage of neurons as they are being edited (Fig. 5a) allowing us to map any root segment used in a query to the closest available snapshot time point (Fig. 5a). This produces an overinclusive set of segments with which we query the materialized database. Additionally, we query the “live” database for all changes to annotations since the used materialization snapshot and add them to the returned set of annotations. The resulting set of annotations is then mapped to the query timestamp using the lineage graph and supervoxel to root lookups, and finally reduced to only include the queried set of root IDs.

The additional logic required to execute arbitrary time point querying introduces an overhead over querying materialized analysis snapshots directly. To quantify this overhead, we turned to the FlyWire dataset for which numerous actively proofread neurons were available. Starting from a materialized snapshot we queried presynapses (Fig. 5c) for individual neurons at several time offsets from the snapshot using the delta query logic (see Methods). The resulting measurements can be categorized in three groups. First, we obtained measurements for presynapse queries of FlyWire neurons that were aligned with a snapshot (median=525 ms, N=121,400, Fig. 5d). Second, we gathered timings for non-snapshot aligned queries where the query neuron did not see any edits since the snapshot version, though its synaptic partners may have (median=978 ms, N=127,775), and, third, where the query neurons were edited since the snapshot (median=1,385 ms, N=12,303).

By comparing measurements from the first two groups for queries to the same neuron, we can obtain the overhead of the additional logic (median=447 ms, Fig. 5e). While query times were well correlated with query size (Ext. Data Fig. 5-1a), we found this offset to be largely constant across queries (Ext. Data Fig. 5-1b). Queries are being slowed down modestly for the third case where the queried segment changed since the last snapshot and an overinclusive query has to be generated (Fig. 5e, f).

### Modular and open design for broad dissemination

We designed CAVE along two broad principles: modularity and openness. Rather than a monolithic application, CAVE is designed as a set of loosely coupled services (Supplemental Table 1). Each CAVE service serves a specific purpose, controls its own data, and is deployed as a docker image to Google Cloud through kubernetes (see Methods). Services can always be added to meet a specific need of a community and replaced with ones that fulfill the same purpose and application interfaces (APIs).

CAVE services can be accessed through authenticated APIs. We developed a Python client (CAVEclient) for programmatic access and adapted the popular viewer neuroglancer^55^ (Fig. 6a) for interactive viewing and editing of the ChunkedGraph segmentation. Further, interactive analysis is enabled through custom dash apps (python-based web apps) that can be extended to serve the needs of any community (Fig. 6a). CAVE’s APIs can be accessed by other tools (Fig. 6a) so long as they authenticate with CAVE’s centralized authentication and authorization server. The analysis package natverse^56^ and the web applications Codex (https://codex.flywire.ai), braincircuits.io (https://braincircuits.io), and NeuVue^57^ already serve as such examples (Fig 6a).

**Figure 6.**
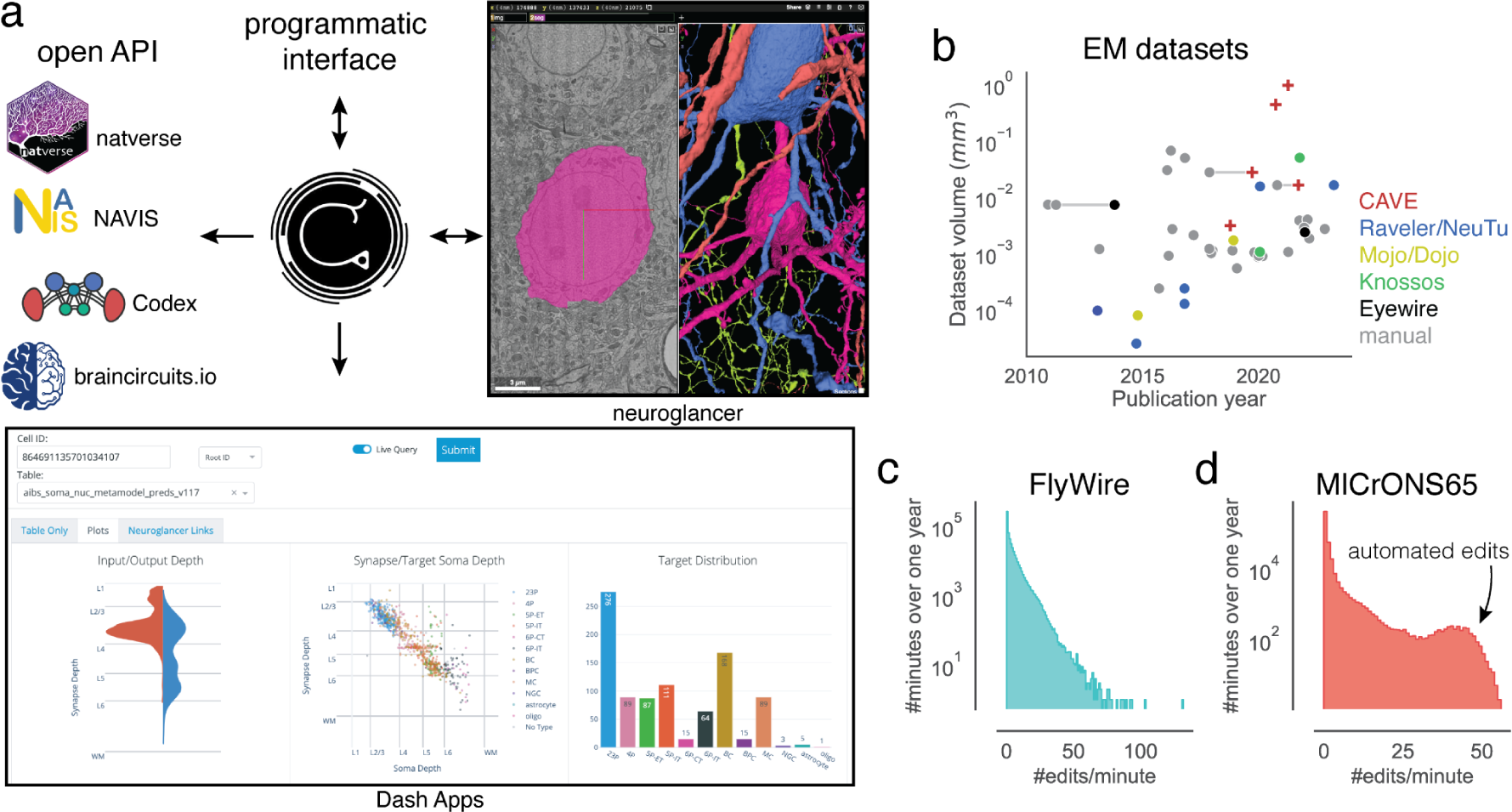
Integration into connectomics projects. (a) CAVE supports multiple interfaces. Besides through programmatic access, users can explore and edit the data in CAVE interactively through neuroglancer or CAVE’s Dash Apps. CAVE integrates with existing and new tools for connectomics though such as natverse^56^, Codex, and braincircuit.io. (b) Datasets published since 2010 by volume and year (volume is plotted on a log-scale). Datasets that were published with manual and semi-automated means are connected with a horizontal gray line. (c) Proofreading rate in edits/min for FlyWire and (d) MICrONS65 over one year of proofreading.

To date, CAVE has facilitated proofreading and analysis of five published datasets with many others in progress (Fig. 6b), including FlyWire^34^, FANC^45^, the MICrONS datasets^17, 18, 33, 49^, and the H01 human dataset^19^. Together, these communities accumulated over four million edits so far by over 500 unique users across the globe. Proofreading by a community puts unpredictable loads onto CAVE. Proofreading rates vary throughout the day with FlyWire seeing as many as >100 edits/minute (Fig. 6c). More than 150,000 edits in MICrONS65 were applied automatically^58^ illustrating how CAVE supports both manual and automated proofreading efforts.

## Discussion

We introduced CAVE, an open-source software infrastructure for managing proofreading, annotations, and analysis by a distributed group of scientists. It is the first system that enables concurrent proofreading and annotation querying at arbitrary timepoints for seamless analysis, and the only system that has successfully demonstrated proofreading of petascale connectomics datasets.

While CAVE demonstrates significant advances, it also combines many features inspired by prior tools for distributed connectome analysis (see ^59^ for a review). CAVE was particularly influenced by CATMAID^40^ which enables collaborative annotation, manual neuron tracing, and analysis, and was used by the *Drosophila* larva community^60^. Similarly, webknossos^42^, Viking^43^, and Knossos^41^ (https://knossos.app) support collaborative manual tracing and annotating. For distributed proofreading, Eyewire^35^ is the closest precedent and was the first tool to distribute block-based proofreading to a community through an interactive browser interface with remeshing capabilities after edits. NeuTu^31^ demonstrated neuron-based proofreading at scale for a restricted group of people that proofread multiple *Drosophila* datasets^61–64^, including the hemibrain^16^.

Analysis of proofread data has so far relied on static exports after proofreading was completed. For instance, NeuPrint^32^ provides analysis of data after it has been proofread in NeuTu and can, in theory, also use exports and materialized snapshots from CAVE. While the reliance on static data is limiting for their use during proofreading, such tools can provide more complex analyses, such as graph queries, through preprocessing of the synapse graph^65^, as illustrated by NeuPrint^32^(https://neuprint.janelia.org), Codex and FlyBrainLab^66^.

Connectomics data, and biological imaging data in general, are being generated at a growing rate. Due to their size, these data are increasingly analyzed by multiple people for a long period of time raising the demand for interoperable and flexible tools that enable simultaneous editing and distributed analysis across multiple user groups. CAVE’s design enables anyone to interface with it to provide specific functionality needed for a given community, providing an example framework for addressing this challenge. This means that others do not need to replicate CAVEs entire functionality, lowering the barrier to entry for the development of new analysis tools. For each new dataset using CAVE, these new analysis tools can automatically be used through CAVE’s common APIs.

In scaling up to petascale datasets, CAVE faced tradeoffs between cost, operational complexity, and performance. In particular, to deploy CAVE a scientific project needs personnel able to manage container based web services, as opposed to standalone desktop tools. Furthermore, we optimized CAVE for large connectomics datasets with dense automated reconstructions and many users. This led us to focus our engineering efforts on making CAVE scale well with respect to cost while maintaining sufficiently high performance, as illustrated by our upgrades to the ChunkedGraph v2. Future scaling to even larger datasets will face similar decisions to reduce costs for proofreading and analysis infrastructure for the same volume. Another significant source for cost is the storage of all annotations in a relational database that can be quickly queried. Automated pipelines now provide accurate and valuable annotations at scale^3, 67^ but storage costs grow linearly with the number of annotations. With the emergence of multi-dimensional annotations demonstrating efficient prediction of semantic information ^68^, new storage solutions will be needed to leverage their power while keeping storage cost in check.

Despite the speedup provided by proofreading of automated segmentations over manual tracing, the manual proofreading component of the reconstruction pipeline remains one of the costliest and slowest steps in the dataset creation process. Further advancements in automated reconstruction will be needed to enable scaling to larger datasets^69^. Instead of improving the automated segmentation directly, tools to automate the proofreading process are emerging^58, 70, 71^. Their application will require new workflows of human-AI interaction^57^. Here, CAVE’s services have already served as a backend system to ingest edits from one automated proofreading pipeline^58^. Due to its broad use, CAVE systems are already holding on to a wealth of data, both annotations and edit histories, that should be leveraged by new automated methods to predict rich annotations of the data and help reduce the need for manual proofreading.

## CAVE related packages

**Supplemental Table 1:**
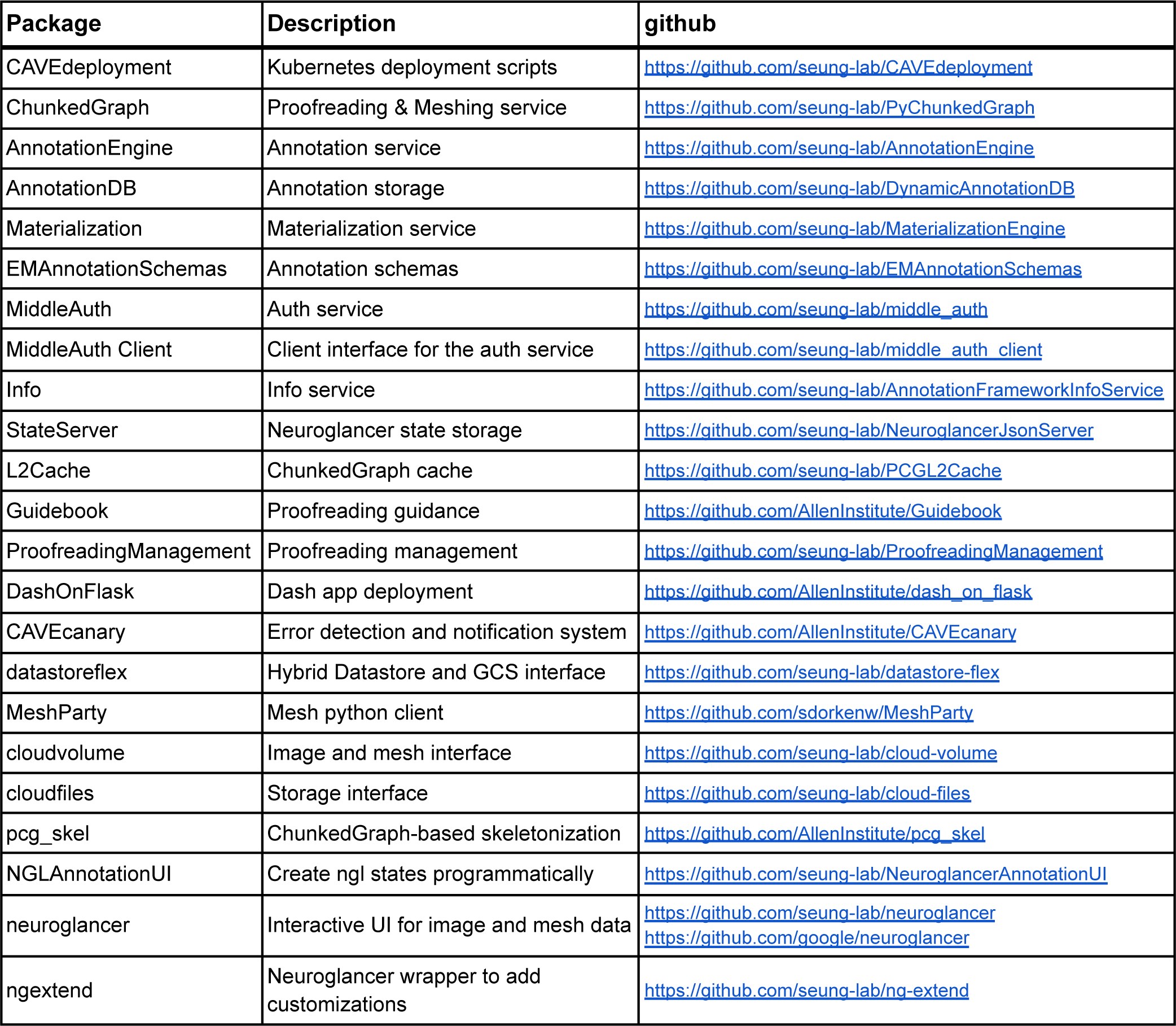
Overview of CAVE packages.

## Acknowledgements

We thank John Wiggins, G. McGrath, and Dave Barlieb for computer system administration and M. Husseini for project administration. We thank Gregory Jefferis, Davi Bock, Eric Perlman, Philipp Schlegel, and Stephan Gerhard for providing feedback on the system design. We thank Gregory Jeffries, Phillip Schlegel, Brock Wester, Stephan Gerhard, Arie Matsliah, Brendan Celli, and Jake Reimer for building tools that leverage the CAVE infrastructure and providing consistent feedback on its performance and implementation. We thank Pedro Nunez Gomez for providing advice about deployment strategies. We would like to thank John Tuthill, Wei-Chung Allen Lee and the FANC community for their collaboration. We would like to thank Viren Jain, and Jeff Lichtman for collaboration on the H01 dataset. We would like to thank Stanley Heinze, Kevin Teodore for collaboration on their datasets. We would like to thank Zetta.ai for collaborations that use CAVE for further datasets. We would like to thank the FlyWire Consortium for collaboration on the FlyWire dataset. We thank the Allen Institute for Brain Science founder, P. G. Allen, for his vision, encouragement and support. Sebastian Seung acknowledges support from the National Institutes of Health (NIH) BRAIN Initiative RF1 MH129268, U24 NS126935, and RF1 MH123400, as well as assistance from Google. Forrest Collman, Nuno da Costa and Clay Reid acknowledge support from NIH RF1MH125932 and from NSF NeuroNex 2 award 2014862. Jakob Troidl and Hanspeter Pfister were supported by NSF grant NCS-FO-2124179. This work was supported by the Intelligence Advanced Research Projects Activity via Department of Interior/Interior Business Center contract numbers D16PC00004, D16PC0005, 2017-17032700004-005 and 2020-20081800401-023. The US Government is authorized to reproduce and distribute reprints for Governmental purposes notwithstanding any copyright annotation thereon. The views and conclusions contained herein are those of the authors and should not be interpreted as necessarily representing the official policies or endorsements, either expressed or implied, of Intelligence Advanced Research Projects Activity, ODNI, Department of Interior/Interior Business Center or the US Government.

## Contributions

SD, FC, CMSM designed CAVE’s core functionalities, and service interactions and layout. SD, AH implemented the ChunkedGraph and the L2 Cache. MC, SD, WS, AH implemented the ChunkedGraph meshing logic. CMSM implemented the improved splitting logic. CJ, NK, JMS, DX extended neuroglancer for proofreading. JMS, VG, JT, HP implemented adapters for and tested CAVE with supervoxel graphs produced by other segmentation pipelines. CJ, SD, FC implemented the authentication system. FC, DB implemented the annotation service and the annotation schema system. DB, FC, SD implemented the annotation database and the materialization service. SD, FC implemented the neuroglancer state server. FC, CMSM, SD, DB implemented the CAVEclient. FC, CMSM, SD implemented MeshParty. CMSM, FC implemented NeuroglancerAnnotationUI. AH, SD implemented datastoreflex. CMSM, FC implemented PCGskel and skeletonization processing. CMSM, FC implemented the dashapps. WS provided support and tools for fast cloud storage access. FC, SD, CMSM, DB, CJ, AH maintained the kubernetes deployments. AH, ALB, BN, CJ, CMSM, DB, DJB, DK, EM, FC, GM, HSS, JAB, JB, JW, KL, KL, LE, MAC, MT, NK, NLT, NMdC, RCR, RL, RT, SCY, SD, SK, SM, SP, SSM, TM, WS, WW, WY, ZJ created the structural MICrONS65 dataset. SD, FC, CMSM wrote the manuscript with contributions from all authors.

## Methods

### Authentication and authorization

The middle-auth service provides a dataset specific authorization layer on top of OAuth2 based authentication. Endpoints allow services to query whether users have different permissions on different service tables. The middle-auth service provides a mapping between service tables and datasets, as well as individual users and groups. Groups then have permissions on datasets.

For example, the ChunkedGraph service has a table named minniev1, so when a user attempts to perform an edit on that table, the service will query a middle-auth endpoint to inquire if that user has “edit” permissions on that table. First, if the user is not logged in, middle-auth will forward them onto Google’s OAuth2 service to authenticate their identity. Upon return, that user is then registered with a unique ID in the middle-auth system. The “minniev1” string is mapped in the “microns” dataset in the ChunkedGraph service namespace, and all the permissions that groups the user is a member of are gathered to see if at least one of them has edit access. If it does, the middle-auth endpoint returns a success, otherwise it returns an unauthorized status code, which is forwarded to the user. This same workflow is used whether or not the user is interacting with the service via python or via neuroglancer.

The programming of this interaction is simplified by the middle-auth-client library, which provides a set of decorators that can be used on flask endpoints to ensure that users are logged in, or that they have particular permissions enabled to access that endpoint. The user’s ID is then made available in the flask global variable dictionary for the service to record which user is performing each request.

### Microservice Architecture

Services are run in docker using a nginx-uwsgi implementation to distribute requests to multiple worker processes operating in a single container. Generally, services have been written in python using the Flask framework, with varying Flask plugins utilized by different services. We use kubernetes to manage container deployment. Kubernetes spins up multiple container pods to increase the number of requests which are handled by each service. Requests are distributed across those pods though load balancing, and an nginx-ingress controller is used to route requests from a single IP to the appropriate service based on the url prefix. Most CAVE services are implemented with a common set of technologies and patterns, though this isn’t strictly a technical requirement. Cert-manager is used in conjunction with CloudDNS to manage and renew SSL certificates.

### Neuroglancer state server

The neuroglancer state server was written as a Python flask app, where the json states are stored in Google Cloud Datastore as a simple key-value store. Keys are state IDs and values are the json encoded neuroglancer state. By passing states through an endpoint, programmatic migration of old state values and formats was possible and enabled seamless changing of user experiences as systems migrated. Posting and retrieving json states are implemented as separate endpoints. In neuroglancer, a “Share” button uses the posting endpoint to upload the json state which returns a state ID. The user is provided with a shortened link that contains a reference to the retrieval endpoint with that state ID as a query argument. We programmed neuroglancer to automatically load states when they are defined in the URL, and so this mechanism effectively allows the reduced size link to be shared easily even when the number of annotations specified in the state is very large.

### ChunkedGraph implementation

The original ChunkedGraph implementation is described elsewhere ^34^ and all concepts described there apply to the ChunkedGraph v2 as well. The ChunkedGraph implements the graphene format which is derived from neuroglancer’s precomputed format used for common segmentations.

### Supervoxel edge storage and retrieval

In the original implementation, all supervoxel edges were stored in BigTable. We devised a new storage scheme where all edges are stored on Google Cloud Storage (GCS). Edges are only accessed for edits. Edges are sorted by chunk and stored as protobufs of compressed arrays. Arrays are compressed using zstandard^72^. We changed the edge reading logic to read edges from a chunk in bulk to minimize access to GCS.

Cross-chunk edges are accessed more often than edges within chunks because they link individual components from subtrees together. They are also used to create an L2 graph, a graph between all L2 chunk components. Because of that, cross chunk edges are also stored in BigTable.

Edges are either “on” or “off” and only the “on” edges contribute to the connected components. The initial ChunkedGraph implementation stored this information alongside the edges. This information was redundant with the ChunkedGraph’s hierarchy. In ChunkedGraph v2 we do not store “on” / “off” information with the edges. We implemented logic to infer the edge state from the ChunkedGraph’s hierarchy directly.

The edge information on GCS is never changed. If new edges need to be inserted, they are stored in BigTable. We call such edges “fake edges” and reserved one row per chunk for them. Every operation reading supervoxel edges from GCS also checks for fake edges. Like all entries in BigTable, fake edges are timestamped allowing for an accurate retrieval of the supervoxel graph for timestamps before their creation.

We implemented new logic to check whether fake edges are needed. When a user commands a merge operation, we first check if there is a path in the local supervoxel graph. If there is, we extract all local edges between the two components that are being merged and process the merge operation with them. If there are no such edges, the original user input is used to insert a fake edge between the two selected supervoxels.

### Common format for all supervoxel graphs

The reimplemented storage of the supervoxel edges does not change and can be used for multiple ChunkedGraphs for the same dataset. This further reduces the per-ChunkedGraph cost. This edge storage also serves as a common format from which the ChunkedGraph can ingest supervoxel graphs created by other segmentation pipelines. In addition to edges, this format also contains component files (protobuf) which store the connected component information inferred by the segmentation pipeline. So far exports to this format were generated for the flood filling segmentation pipeline ^26^ and the LSD segmentation pipeline ^27^.

### ChunkedGraph Meshing

#### Meshing procedure

We use zmesh (https://github.com/seung-lab/zmesh) for meshing of the segmentation. Every L2 chunk component is meshed at MIP 2 and the mesh stored on GCS. Some L2 components only consist of a few pixels and might not be meshed at all. L2 meshes are then stitched on the layers above following the ChunkedGraph’s hierarchy up to a stop layer (e.g. 6) at which many meshes are too large to be held in the memory of a worker node or time constraints of queueing systems (e.g. Amazon SQS) are surpassed.

#### Mesh storage

The large number of L2 IDs (>10^9^) translates into a large number of L2 meshes which would be expensive to store as individual files on GCS because of the high cost of write operations (e.g., GCS charges $0.005/1,000 write operations). For the initial meshing of all segments in the ChunkedGraph after ingest, we store meshes in sharded files. For each L2 chunk, we store all meshes in one sharded file using cloudfiles (https://github.com/seung-lab/cloud-files). This pattern is repeated for higher layers, where stitched versions of meshes are stored, reducing the number of files that need to be downloaded for any given neuron. The sharded format allows retrieval of byte ranges from each shard but adds two additional reads to the header for retrieving the byte range. Note, this is a different arrangement from the original precomputed sharded mesh format, where all the mesh fragments from a single neuron can be found in a shard.

Sharded files cannot be extended, only rewritten. This posed a problem for meshes that were created for new components after an edit. However, these are few in number in comparison to the number of meshes from the initial meshing run. Consequently, we store each of these newly generated mesh fragments as a single file on GCS. We refer to this format as “hybrid mesh format” because it uses both sharded and single file storage.

Each mesh is compressed using the Draco format for which we wrote and maintain a Python client (https://github.com/seung-lab/DracoPy). The Draco format is a lossy mesh compression format where every mesh vertex is moved to the closest grid node. The grid’s spacing determines the compression factor. We place a global grid onto the dataset such that meshes retrieved from different chunks can be merged through overlapping vertices.

#### Mesh retrieval

Neuroglancer’s precomputed format requires a manifest per segment outlining the mesh fragments that need to be read from GCS to produce a complete rendering of a segment. Every edit creates a new cell segment ID with a new manifest. Instead of precalculating and storing all manifest, the ChunkedGraph produces manifest for segments on the fly from the hierarchy of a neuron. Using the timestamp of the fragment ID, the ChunkedGraph can determine whether the fragment is stored in sharded or file-based storage and provide instructions to accordingly.

Cloudvolume implements all necessary interactions with the ChunkedGraph and can be used to programmatically read meshes. MeshParty wraps this functionality and adds convenience functionality such as caching of meshes, on-disk and in-memory, and provides further capabilities such as mesh rendering using VTK.

### L2-Cache

Features for each L2 ID are calculate on the binarized segmentation. For each L2 ID, we currently calculate the following features:

Representative coordinate, volume, area, principal components, mean and maximum value of the euclidean distance transform, number of voxels at each chunk boundary intersection.

Area calculations are difficult to perform and are easily inflated by rough surfaces. However, smoothed measurements are ill-defined and expensive to obtain. Thus, our area measurements overestimate the actual area of a neuron. We calculate areas by shifting the segmentation in each dimension and finding all voxels where the segment of interest overlaps with other segments. We count up the surfaces and adjust for resolution. L2 features are stored in BigTable. Every L2 ID matches to a row in BigTable and contains a column for each feature.

### L2 skeletonization

Skeletons were generated using a graph-based generalization of the TEASAR algorithm ^53^ using L2 chunks. For a given root id, we query the ChunkedGraph for its component L2 ids and the list of which L2 chunks are directly adjacent with which others, either via supervoxels that spanned chunk boundaries or proofreading edits that introduced edges between chunks. We next query the L2-Cache to identify the representative coordinate (and other properties) for each L2 id and use this information to generate a graph where each vertex is a single L2 chunk and edges have a weight given by the distance between representative coordinates of adjacent chunks. Following the TEASAR algorithm, we identify a root node (for example, the closest vertex to a cell body centroid or the base of the axon from a peripheral sensory neuron) and find the most distant vertex on the graph. The vertices along the shortest path from the distant vertex to the root node are assigned to the skeleton and we “invalidate” vertices within a distance parameter provided by the user. Importantly, we store a mapping from each invalidated graph vertex to the closest skeleton vertex. We iterate this process, using the most distant un-invalidated vertex and the shortest path to the existing skeleton, until all vertices are invalidated.

To associate synapses with vertices on the skeleton, we get the supervoxel id of the bound spatial point(s) associated with the annotation and use the ChunkedGraph to look up its associated L2 id(s). We then assign synapses to graph vertices via L2 id and use the invalidation mapping to associate graph vertices to supervoxel vertices. Similarly, L2 properties such as volume or surface area for regions of a skeleton can be computed by summing the appropriate values from the L2 Cache via the associated graph vertices. The core skeletonization process was implemented in MeshParty and the interaction with the ChunkedGraph is handled through the python library “pcg_skel”.

### Schemas implementation

Annotation schemas were defined in python code, where they are constructed using the Marshmallow library. Each schema contains at least one field of the custom class “BoundSpatialPoint”. This field implicitly creates fields for positions, supervoxel and their associated root ids. The annotation and materialization process can thus also dynamically locate BoundSpatialPoint fields and use them to execute the generic workflow of supervoxel and root id lookup described in the materialization process.

Reference annotations were defined as a custom Schema subclass with a target id field associated with them. Postgres data access and storage was facilitated by a module which automatically constructed SQLalchemy models, using GeoAlchemy to describe spatial positions as spatially indexed 3D points. This model creation code automatically adds spatial indices to spatial points, and SQL indices to associated root id columns to facilitate fast querying. It also generates the foreign key constraints associated with reference annotations. Each schema is assigned a unique string for identification, and is used by libraries to indicate what schema a table utilizes. Reference annotations may or may not have their own set of BoundSpatialPoints, and their model creation requires an extra parameter to create a foreign key between the target id column of the reference table and the id column of the table that is being referenced.

The EMAnnotationSchemas repository is the source of truth for what kinds of schemas can be initialized, and the community can contribute suggestions through pull requests to this library. Because all model creation code is written generically, extending the schemas supported is easy. This code is then used both as a library in other services, but also as a flask based web service which makes a dynamic list of schemas and their structure as jsonschema, facilitated by the marshmallow-jsonschema library, available.

### Annotation Service

The annotation service manages the creation of new annotation tables as well as creation, deletion and updating of annotations within tables. Annotation tables are stored in Google Cloud SQL using PostgreSQL through a library called DynamicAnnotationDB. Annotation tables of any schema can be created and multiple tables with the same schema may exist. When creating a table users provide metadata about the table via a REST endpoint. These include a description, read and write permissions, and the resolution of the spatial points. The permission model currently allows for 3 levels of permission for both read and write. “PRIVATE” allows only that user to read or write, “GROUP” allows for users in the same group (see authorization) to read or write, “PUBLIC” allows for all users with read or write permissions on the dataset to read or write that table. The default permissions are “PRIVATE” write but “PUBLIC” read to encourage data sharing and reuse within communities.

If the user is creating a table with a reference schema, then they also must specify the name of the table that is being referenced. The service then utilizes the DynamicAnnotationDB library to create the table within the live SQL database and stores the metadata about the table in a separate metadata table. Annotations can be posted through a separate endpoint which accepts json serialized versions of annotations. Annotations are then validated against the schema using marshmallow and the SQLalchemy model is dynamically generated by the schema library. Annotations are then inserted into the PostgreSQL database, after associating a creation timestamp to the annotation.

Upon insertion, the annotation service sends a notification to the Materialization service to trigger supervoxel lookups for the recently added annotations. Deletion is implemented virtually by marking the timestamp of deletion in order to enable point in time consistent querying.

Updates are represented as a combination of remove and add operations. The CAVEclient has python functions for facilitating client side validation and packaging of annotations for the REST endpoint, including support for processing pandas dataframes. In addition to the API, the service provides a human readable website interface for browsing existing tables.

### Materialization implementation

The Materialization Engine updates segmentation data and creates databases that combine spatial annotation points and segmentation information. There are two types of databases that the system uses, one database for the “live” dataset and multiple for materialized snapshots. The live database is the one written to by the Annotation Service, and actively managed by the Materialization service to keep root ids coherent and up to date for all BoundSpatialPoints in all tables. The frozen databases are copies of a time-locked state of the live database’s segmentation and annotation information, used to facilitate consistent querying. To keep the data in sync the backend leverages Celery, a python based distributed task queue, that allows for scaling and distributing parallel workloads. Using dynamic Celery-based workflows, the Materialization Engine runs periodic tasks to keep the segmentation information up to date from the proofreading efforts and provides copies of “frozen” datasets at a fixed interval for analysis.

The Materialization Engine is deployed to a kubernetes cluster where celery is run on pods. Two types of celery pods are deployed for CAVE: producers and consumers. Producers create workflows that dynamically generate tasks that the consumer pods will subsequently execute.

### Annotation query implementation

The Materialization service also provides a query API for users to query both materialized versions of the database, as well an endpoint which implements a workflow that enables arbitrary moment in time querying of the data, with nearly identical features. Both sets of endpoints enable arbitrary filters on columns from the annotation tables, with inclusive, exclusive and strictly equal filter options, as well as bounding box spatial queries on all spatial points. By filtering on the segment id columns of associated BoundSpatialPoints, users can efficiently extract all annotations for individual cells. For example, this lets users retrieve all input or output synapses from a particular set of neurons, or allows users to query what cell type annotations are associated with a particular set of cells. A join query endpoint allows users to create arbitrary inner join queries on annotation tables, with the same filter criteria. Queries return data either as PyArrow binary dataframes, which is faster and more efficient, or as json serialized objects, which is more cross platform compatible, depending on a query parameter option. To prevent queries from accidently requesting multiple gigabytes of data, an arbitrary configurable upper bound on the number of rows that are requested from the SQL database is enforced. Presently our deployed systems have configured this to be 500,000 rows, though users can distribute more data by executing multiple requests in parallel.

Although the live query endpoint appears similar to the materialized endpoint to the user, the workflow in the background is more complex. In addition to the filters described above, users must specify a timestamp they are interested in querying for a live query. First, the system uses the ChunkedGraph’s lineage graph to translate all the filter parameters referencing segment IDs into an over-inclusive set of related segment IDs that are present in the closest materialized timepoint. Equality filters are translated to inclusive filters in this process. This translated query is then executed against the materialized database to retrieve all the annotations that are potentially related to the users query. Then, for all the segment IDs that are not valid at the user provided timestamp, the ChunkedGraph API is queried using the associated supervoxel ids to determine the correct segment ID for those BoundSpatialPoints. This covers any changes that might have happened in the segmentation data between materialization and the queried timepoint, but does not account for changes in the annotation data that might have happened in that interval. Therefore, a second query is executed on the “live” database using a filter on the created and deleted columns to extract any annotation rows that were added or removed on the queried tables. Note, if the closest materialization point is in fact in the future, then the meaning of addition and removal is inverted with respect to this step. The annotation service also tracks a timestamp for when a table was last modified to skip this step if there is no possibility the table was altered in the interval between materialization and the query. Filters on segment IDs must be ignored in this process because there can be no guarantee of consistency on the live database due to ongoing and distributed update operations. Once these new and deleted annotation rows are retrieved, the same process of updating expired segment IDs using the ChunkedGraph API is applied. Rows from the materialized query which exist as deletions in the live database query are removed, and rows which were added are concatenated to the result.

Finally, the original query filters on segment IDs are applied to these aggregated results to remove any annotations which are not relevant to the users query.

### ChunkedGraph performance measurements

We measured server response times for all endpoints served by the ChunkedGraph from all users for several weeks while proofreading was progressing as normal. These numbers reflect real interactions and are affected by server and database load and are therefore an underestimate of the capability of our system. The most common requests are root to leave requests as they are executed every time a user moves their field of view in neuroglancer. We sampled a random set of these interactions. For all others, we sampled all interactions.

### Morphological feature performance measurements

We used a compute node on Google Cloud to execute programmatic queries to CAVE. First, we selected representative sets of neurons from each dataset (FlyWire, MICrONS65): We used neurons from MICrONS65 that were included in a recent circuit analysis^73^ representing most proofread neurons in the dataset, and all neurons that were marked as proofread from FlyWire. Then we randomly sampled neurons from each list and queried: (1) All L2 IDs from the ChunkedGraph, (2) All volume measurements from the L2 Cache for these L2 IDs. We finally added up all volume measurements for a total volume. We processed neurons sequentially for multiple days and recorded all measurements. We average time measurements for neurons for which we gathered multiple measurements. Measured performances were affected by the current load on the system.

### Annotation query performance measurements

We used the proofread neurons from FlyWire for this analysis. Starting from materialization 571, we executed presynapse queries at time offsets of 0, 1, 10, 20, 40, 100, 400, 800 hours recreating realistic queries. It should be noted that most queries are within 24 h of a materialization version. We precomputed the neuron IDs and number of edits after the materialization version for those timepoints and created a list of tuples containing (segment id, timestamp) from which we randomly sampled entries and executed a presynapse queries.

Queries were executed sequentially.

